# Early farmers from across Europe directly descended from Neolithic Aegeans

**DOI:** 10.1101/032763

**Authors:** Zuzana Hofmanová, Susanne Kreutzer, Garrett Hellenthal, Christian Sell, Yoan Diekmann, David Díez-del-Molino, Lucy van Dorp, Saioa López, Athanasios Kousathanas, Vivian Link, Karola Kirsanow, Lara M. Cassidy, Rui Martiniano, Melanie Strobel, Amelie Scheu, Kostas Kotsakis, Paul Halstead, Sevi Triantaphyllou, Nina Kyparissi-Apostolika, Dushanka-Christina Urem-Kotsou, Christina Ziota, Fotini Adaktylou, Shyamalika Gopalan, Dean M. Bobo, Laura Winkelbach, Jens Blöcher, Martina Unterländer, Christoph Leuenberger, Çiler Çilingiroğlu, Barbara Horejs, Fokke Gerritsen, Stephen Shennan, Daniel G. Bradley, Mathias Currat, Krishna R. Veeramah, Daniel Wegmann, Mark G. Thomas, Christina Papageorgopoulou, Joachim Burger

## Abstract

Farming and sedentism first appear in southwest Asia during the early Holocene and later spread to neighboring regions, including Europe, along multiple dispersal routes. Conspicuous uncertainties remain about the relative roles of migration, cultural diffusion and admixture with local foragers in the early Neolithisation of Europe. Here we present paleogenomic data for five Neolithic individuals from northwestern Turkey and northern Greece – spanning the time and region of the earliest spread of farming into Europe. We observe striking genetic similarity both among Aegean early farmers and with those from across Europe. Our study demonstrates a direct genetic link between Mediterranean and Central European early farmers and those of Greece and Anatolia, extending the European Neolithic migratory chain all the way back to southwestern Asia.

---

It is well established that farming was introduced to Europe from Anatolia, but the extent to which its spread was mediated by demic expansion of Anatolian farmers, or by the transmission of farming technologies and lifeways to indigenous hunter-gatherers without a major concomitant migration of people, has been the subject of considerable debate. Paleogenetic studies (*1–6*) of late hunter-gatherers and early farmers indicate a dominant role of migration in the transition to farming in central and northern Europe, with evidence of only limited hunter-gatherer admixture into early Neolithic populations. However, the exact origin of central European early farmers, in the Balkans, Greece or Anatolia remains an open question.

Recent radiocarbon dating indicates that by 6,600 to 6,500 cal BCE, sedentary farming communities were established in northwest Anatolia, at sites such as Barcın, Menteşe, and Aktopraklık C (*7, 8*), and in coastal west Anatolia, at sites like Çukuriçi and Ulucak (*9, 10*) (Fig. 1), but did not expand north of the Aegean for another several hundred years (*11*). All these sites show material culture affinities with the central and southwest Anatolian Neolithic (*12, 13*). Early Greek Neolithic sites, such as Franchthi Cave in the Peloponnese (*14*), Knossos in Crete (*15*) and Mauropigi, Paliambela and Revenia in northern Greece (*16–18*) date to a similar period. The distribution of obsidian from the Cycladic islands, as well as similarities in material culture, suggest extensive interactions since the Mesolithic and a coeval Neolithic on both sides of the Aegean (*17*). While it has been argued that *in situ* Aegean Mesolithic hunter-gatherers played a major role in the ‘Neolithisation’ of Greece (*14*), the presence of domesticated forms of plants and animals is a good indication of extra-local Neolithic dispersals into the area (*19*).

**Figure 1:**
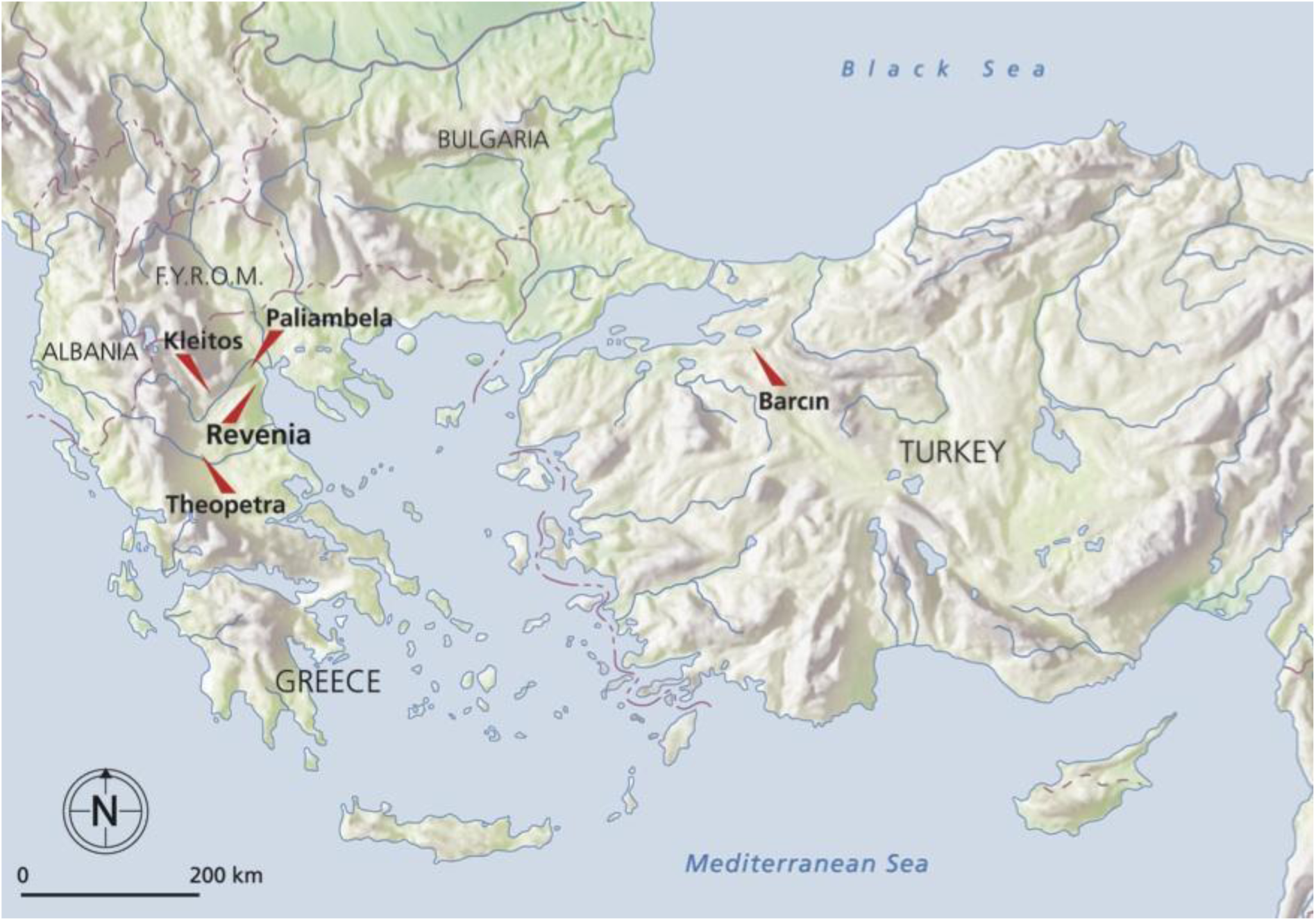
North Aegean archaeological sites investigated in Turkey and Greece.

We present five ancient genomes from the European and Asian sides of the northern Aegean (Fig. 1); three sequenced to relatively high coverage (~3–8x) enabling diploid calls using a novel SNP calling method that accurately accounts for post-mortem damage. Two of the higher coverage genomes are from Barcın, south of the Marmara Sea in Turkey, one of the earliest Neolithic sites in northwestern Anatolia (Bar8 and Bar31, Table 1). On the European side of the Aegean, one genome is from the early Neolithic site of Revenia (Rev5), and the remaining two are from the late and final Neolithic sites of Paliambela (Pal7) and Kleitos (Klei10) dating approximately 2,000 years later (Table 1).

**Table 1:**
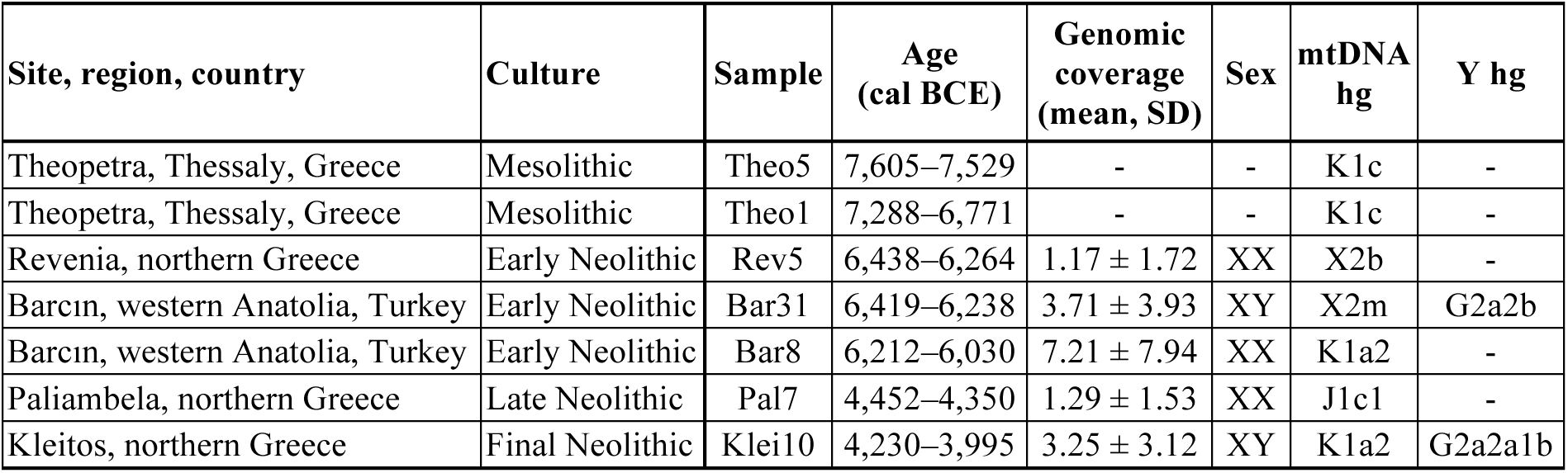
Neolithic and Mesolithic samples analysed.

| Site, region, country | Culture | Sample | Age (cal BCE) | Genomic coverage (mean, SD) | Sex | mtDNA hg | Y hg |
| --- | --- | --- | --- | --- | --- | --- | --- |
| Theopetra, Thessaly, Greece | Mesolithic | Theo5 | 7,605–7,529 | - | - | K1c | - |
| Theopetra, Thessaly, Greece | Mesolithic | Theo1 | 7,288–6,771 | - | - | K1c | - |
| Revenia, northern Greece | Early Neolithic | Rev5 | 6,438–6,264 | $1.17 \pm 1.72$ | XX | X2b | - |
| Barcin, western Anatolia, Turkey | Early Neolithic | Bar31 | 6,419–6,238 | $3.71 \pm 3.93$ | XY | X2m | G2a2b |
| Barcin, western Anatolia, Turkey | Early Neolithic | Bar8 | 6,212–6,030 | $7.21 \pm 7.94$ | XX | K1a2 | - |
| Paliambela, northern Greece | Late Neolithic | Pal7 | 4,452–4,350 | $1.29 \pm 1.53$ | XX | J1c1 | - |
| Kleitos, northern Greece | Final Neolithic | Klei10 | 4,230–3,995 | $3.25 \pm 3.12$ | XY | K1a2 | G2a2a1b |

Dates calibrated using Oxcal v4.2.2 and the Intcal13 calibration curve. For details on ^14^C dating see Supplementary Information, section 1.

The mtDNA haplogroups of all five individuals are typical of those found in central European Neolithic farmers and modern Europeans, but not of European Mesolithic hunter-gatherers. Likewise, the Y-chromosomes of the two male individuals belong to haplogroup G2a2, which has been observed in European Neolithic farmers (*3, 20, 21*), Ötzi, the Tyrolean Iceman (*22*), and modern western and southwestern Eurasian populations, but not in any pre-Neolithic European hunter-gatherers. However, the mitochondrial haplogroups of two additional less well-preserved Greek Mesolithic individuals (Theo1, Theo5) belong to lineages observed in Neolithic farmers from across Europe, consistent with Aegean and possibly central Anatolian Neolithic populations, unlike central European Neolithic populations, being the direct descendants of the preceding Mesolithic peoples that inhabited broadly the same region. Two recently published pre-Neolithic genomes from the Caucasus (*23*) appear to be highly differentiated from the genomes presented here and most likely represent a forager population distinct from the Epipaleolithic/Mesolithic precursors of the early Aegean farmers (Fig. 2).

**Figure 2:**
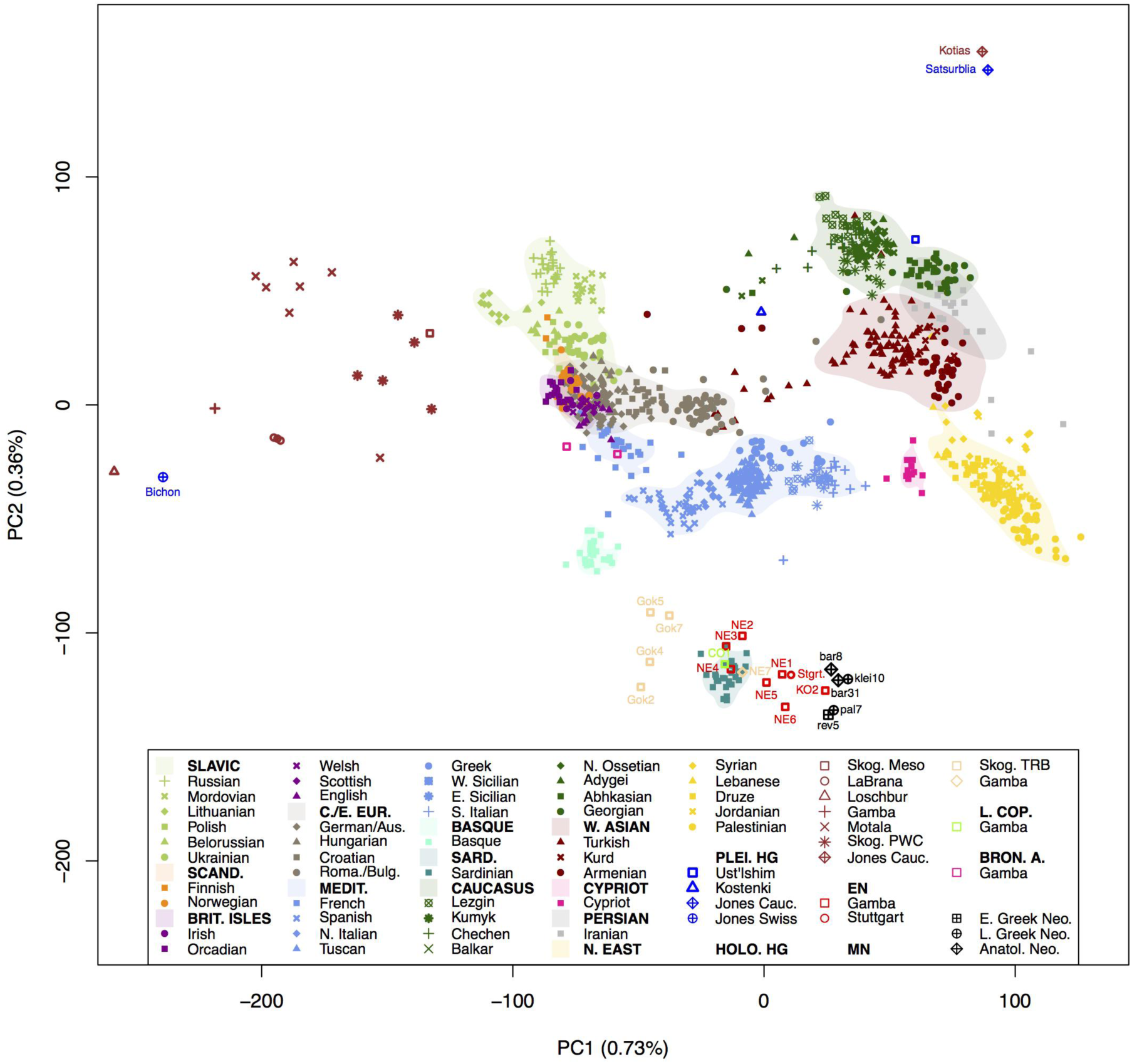
PCA of modern reference populations and projected ancient individuals. The Greek and Anatolian samples reported here cluster tightly with other European farmers close to modern-day Sardinians, however, are clearly distinct from previously published Caucasian Hunter-Gatherers (*23*). This excludes the latter as potential ancestral source population for early European farmers and suggests strong genetic structure in hunter-gatherers of southwest Asia.

The first two dimensions of variation from principal components analysis (PCA) reveal a tight clustering of all five Aegean Neolithic genomes with Early Neolithic genomes from central and southern Europe (*2, 3, 24*) (Fig. 2).

To examine this clustering of Early Neolithic farmers in more detail, we calculated outgroup *f*3 statistics (*25*) of the form *f*3 (Khomani; TEST, Greek/Anatolian), where TEST is one of the available ancient European genomes; ≠Khomani San were selected as an outgroup as they are considered to be the most diverged extant human population. Consistent with their PCA clustering, the northern Aegean genomes share high levels of genetic drift amongst each other, and with all other previously characterized European Neolithic genomes, including early Neolithic from northern Spain, Hungary and central Europe. Given the archaeological context of the different samples, the most parsimonious explanation for this shared drift is migration of early European farmers from the northern Aegean into and across Europe.

To better characterize this inferred migration, we modeled each ancient genome as a mixture of DNA from other ancient and/or modern genomes (*26, 27*) (Fig. 3). Under this framework the oldest Anatolian genome (Bar31) was inferred to contribute the highest amount of genetic ancestry (30–50%) to the Early Neolithic genomes from Greece (Rev5), Hungary (*24*) and Germany (*2*) compared to any other ancient or modern samples, with the next highest contributors being other ancient Greek (Pal7, Klei10) and/or Anatolian (Bar8) genome. In contrast, contributions from the Hungarian and German Neolithic genomes to any of the Anatolian or Greek ancient genomes were consistently smaller (<11%). Such an asymmetric pattern is indicative of founder effects (*28*) in Hungary, Germany, and possibly Greece from a source that appears to be most genetically similar to Bar31. Consistent with this, we found fewer short runs of homozygosity (ROH< 1.6Mb) in our high coverage Anatolian sample (Bar8) than in Early Neolithic genomes from Germany and Hungary. However, while these results conform to a Neolithic dispersal from Anatolia to Greece, and then to the rest of Europe, it is not possible to infer a direction for dispersal within the Aegean with statistical confidence since both the Greek and Anatolian genomes copy from each other to a similar extent. We therefore see the origins of European farmers equally well represented by Early Neolithic Greek and northwestern Anatolian genomes (*29*).

**Figure 3:**
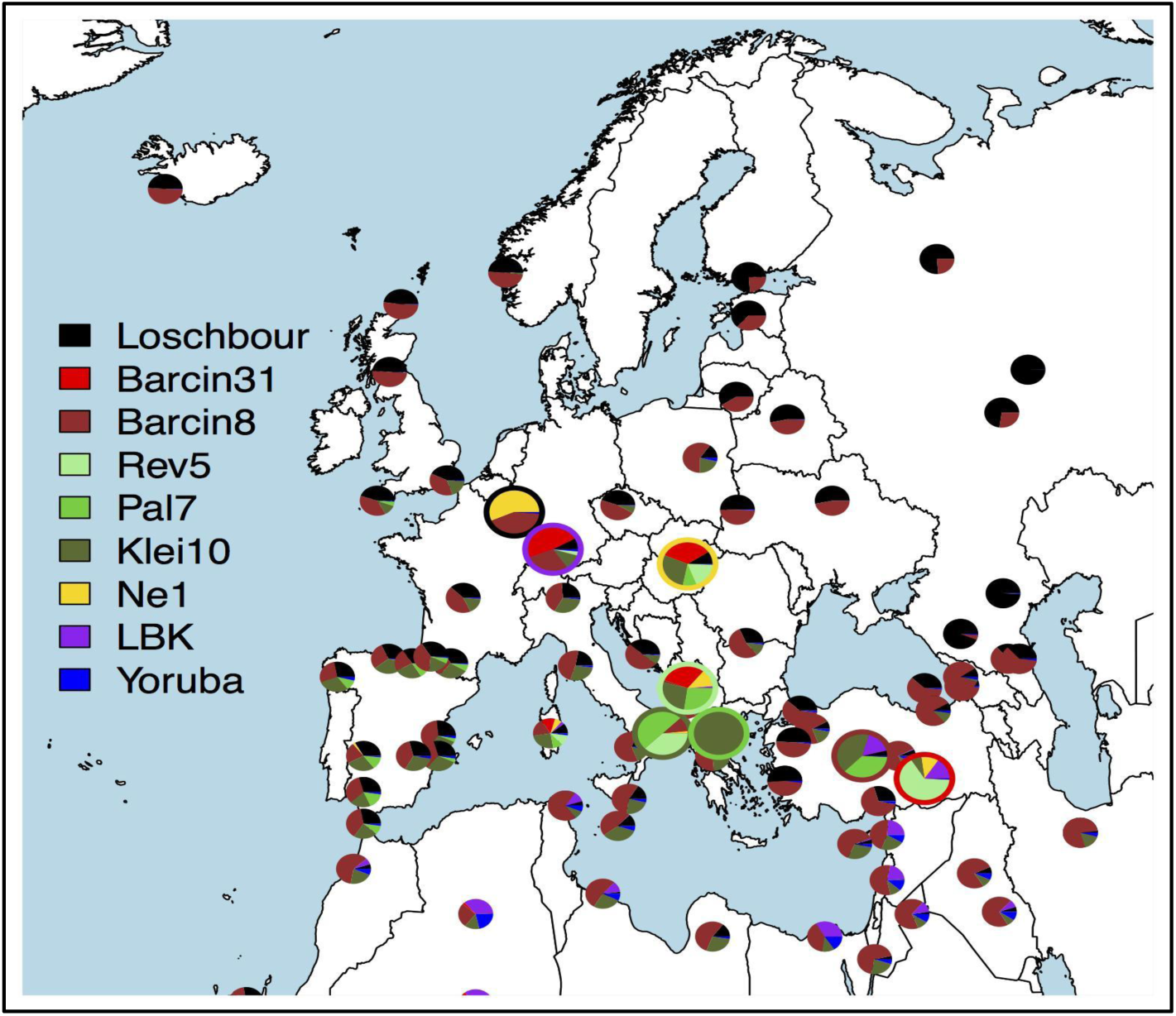
Inferred mixture coefficients when forming each modern (small pies) and ancient (large pies, enclosed by borders matching key at left) group as a mixture of the modern-day Yoruba from Africa and the ancient samples shown in the key at left.

It is widely believed that farming spread into Europe along both Mediterranean and central European routes, but the extent to which this process involved demic dispersals from the Aegean has long been a matter of debate (*30*). We applied *f4* statistics to examine whether the Spanish Neolithic farmers shared more drift with the Early Neolithic genome from Germany than with the Aegeans, which would be expected if Neolithic populations first reached southwestern Europe via central Europe. We found no support for this hypothesis: none of the early farmers in Europe shared significantly more drift with one another than with Aegean farmers. This result is consistent with early farmers migrating from the Aegean via at least two independent routes into central and southwestern Europe, and with inferences made on the basis of archaeological evidence (*31, 32*).

Given the Aegean is the likely origin of European Neolithic farmers, we utilized Bar8 and Bar31 as putative sources to assess the extent of hunter-gatherer admixture in European farmers through the Neolithic. *f*4 statistics of the form *f*4(Neolithic farmer, Anatolian, HG, Khomani) indicated small but significant amounts of hunter-gatherer admixture (at least in comparison to Anatolians) into both Spanish and Hungarian early farmer genomes, and interestingly, the Early Neolithic Greek genome. Our mixture modeling analysis also inferred small genetic contributions from the Loschbour hunter-gatherer genome (2–10%) to each of the Early Neolithic Hungarian and German genomes, but little evidence of contributions to any Aegean genomes. These results suggest that mixing between migrating farmers and local hunter-gatherers occurred sporadically throughout the continent even in the earliest stages of the Neolithic, but only at low levels. However, consistent with previous findings (*3*), both *f4* statistics and ADMIXTURE analysis indicate a substantial increase in hunter-gatherer ancestry transitioning into the Middle Neolithic across Europe, while Late Neolithic farmers also demonstrate a considerable input of ancestry from steppe populations.

Modern Anatolian and Aegean populations do not appear to be the direct descendents of Neolithic peoples from the same region. Indeed, our mixture model comparison of the Aegean genomes to >200 modern groups^2^ indicates low affinity between the two Anatolian Neolithic genomes and seven of eight modern Turkish samples (the eighth is from Trabzon on the Black Sea coast, a long-standing area of Pontic Greek settlement). Furthermore, when we form each Anatolian Neolithic genome as a mixture of all modern groups, we infer no contributions from groups in southeastern Anatolia and the Levant where the earliest Neolithic sites are found. Similarly, comparison of allele sharing between ancient and modern genomes to those expected under population continuity indicates Neolithic to modern discontinuity in Greece and western Anatolia, unless ancestral populations were unrealistically small. Instead, our mixing analysis shows that each Aegean Neolithic genome closely corresponds genetically to modern Mediterraneans, and in particular Sardinians (as also seen in the PCA and outgroup *f3* statistics), with few substantial contributions from elsewhere.

Over the last 6 years ancient DNA studies have transformed our understanding of the European Neolithic transition (*1–6, 24, 29*), demonstrating a crucial role for migration in central and southwestern Europe. Our results bookend this transformative understanding by extending the unbroken trail of ancestry and migration all the way back to southwestern Asia. The lack of shared drift among central and southwestern Early Neolithic farmers to the exclusion of the genomes presented here suggests that Aegean Neolithic populations can be considered the root for all early European farmers and their colonization routes. A key remaining question is whether this unbroken trail of ancestry and migration extends all the way back to southeastern Anatolia and the Fertile Crescent, where the earliest Neolithic sites in the world are found. Regardless of whether the Aegean early farmers were ultimately descended from western or central Anatolian, or even Levantine hunter-gatherer, the differences between the ancient genomes presented here and those from the Caucasus (*23*) indicates that there was considerable structuring of forager populations in southwest Asia prior to the transition to farming.

The dissimilarity and lack of continuity of the Early Neolithic Aegean genomes to modern Turkish and Levantine populations, in contrast to those of early central and southwestern European farmer and modern Mediterraneans, is best explained by subsequent gene-flow into Anatolia from yet unknown sources.

## Acknowledgments

ZH and RM are supported by a Marie Curie Initial Training Network (BEAN / Bridging the European and Anatolian Neolithic, GA No: 289966) awarded to MC, SS, DGB, MGT, and JB. CP, JB, and SK received funding from DFG (BU 1403/6–1). CP and JB received funding from the Alexander von Humboldt Foundation. CS and MS were supported by the EU: SYNTHESYS / Synthesis of Systematic Resources, GA No: 226506-CP-CSA-INFRA, DFG: (BO 4119/1) and Volkswagenstiftung (FKZ: 87161). AS was supported by the EU: CodeX Project No: 295729. KoKo, ST, DCUK, PH, CP were co-financed by the EU and Greek national funds (NSRF). CP, MU, KoKo, ST, DCUK were co-financed by the EU Social Fund and the Greek national funds, program: ARISTEIA II. MC was supported by Swiss NSF grant 31003A_156853. AK, DW were supported by Swiss NSF grant 31003A_149920. SL is supported by BBSRC (Grant Number BB/L009382/1). LvD is supported by CoMPLEX via EPSRC (Grant Number EP/F500351/1). GH is supported by a Sir Henry Dale Fellowship jointly funded by the Wellcome Trust and the Royal Society (Grant Number 098386/Z/12/Z) and supported by the National Institute for Health Research University College London Hospitals Biomedical Research Centre. MGT and YD are supported by a Wellcome Trust Senior Research Fellowship awarded to MGT. JB is grateful for support by the University of Mainz and the HPC cluster MOGON (funded by DFG; INST 247/602–1 FUGG). We thank Songül Alpaslan for help with sampling in Barcın and Eleni Stravopodi for help with sampling in Theopetra.

Accession numbers: Mitochondrial genome sequences are deposited in GenBank (KU171094-KU171100). Genomic data are available at ENA with the accession number PRJEB11848 in BAM format.

